# A Large-Scale Investigation of White Matter Microstructural Associations with Reading Ability

**DOI:** 10.1101/2021.08.26.456137

**Authors:** Steven L. Meisler, John D.E. Gabrieli

## Abstract

Reading involves the functioning of a widely distributed brain network, and white matter tracts are responsible for transmitting information between constituent network nodes. Several studies have analyzed fiber bundle microstructural properties to shed insights into the neural basis of reading abilities and disabilities. Findings have been inconsistent, potentially due to small sample sizes and varying methodology. To address this, we analyzed a large data set of 686 children ages 5-18 using state-of-the-art neuroimaging acquisitions and processing techniques. We searched for associations between fractional anisotropy (FA) and single-word and single-nonword reading skills in children with diverse reading abilities across multiple tracts previously thought to contribute to reading. We also looked for group differences in tract FA between typically reading children and children with reading disabilities. FA of the white matter increased with age across all participants. There were no significant correlations between overall reading abilities and tract FAs across all children, and no significant group differences in tract FA between children with and without reading disabilities. There were associations between FA and nonword reading ability in older children (ages 9 and above). Higher FA in the right superior longitudinal fasciculus (SLF) and left inferior cerebellar peduncle (ICP) correlated with better nonword reading skills. These results suggest that letter-sound correspondence skills, as measured by nonword reading, are associated with greater white matter coherence among older children in these two tracts, as indexed by higher FA.

## 1 Introduction

The development of reading skills is vital for progress in education and communication. Reading disability is the most prevalent learning disability and is characterized by difficulty with word reading accuracy and/or fluency (Roongpraiwan et al., 2002; Lyon et al., 2003). Multiple studies have reported structural and functional brain differences in children and adults with reading disability (Pugh et al., 2000; Eckert, 2004; Maisog et al., 2008; Linkersdörfer et al., 2012; Cattinelli et al., 2013; Richlan et al., 2013; Jednorog et al., 2015), including differences in white matter pathways as measured by diffusion tensor imaging (DTI) (Vandermosten et al., 2012b). Studies of this nature often report the relationship between reading scores and tract fractional anisotropy (FA), which quantifies how direction-dependent water movement is in a given area (Basser and Pierpaoli, 1996; Hagmann et al., 2006) and has been interpreted as an index of white matter coherence (Beaulieu, 2002). Such associations may be important to identify because these fiber bundles of myelinated axons connect the gray matter regions that support reading, and especially the left-hemisphere reading network comprised of the angular gyrus, precuneus, middle temporal gyrus, superior temporal gyrus (including Wernicke’s area), fusiform gyrus (including the visual word form area), and inferior frontal gyrus (including Broca’s area) (Cattinelli et al., 2013; Wandell and Yeatman, 2013; Murphy et al., 2019).

Studies of white matter microstructural properties as they relate to reading abilities in children and adults have yielded interesting, albeit sometimes inconsistent, results. The most commonly implicated tract in DTI studies of reading skills is the superior longitudinal fasciculus (SLF), particularly in the left hemisphere, which connects frontal and temporoparietal brain regions (Wang et al., 2016). Multiple studies have found that higher FA in left and/or right SLF was associated with better reading outcomes, whether that manifested from group comparisons between dyslexic and typically reading individuals (Richards et al., 2008; Steinbrink et al., 2008; Carter et al., 2009; Marino et al., 2014) or correlations with reading test scores on a continuous scale (Steinbrink et al., 2008; Carter et al., 2009; Hoeft et al., 2011; Feldman et al., 2012; Lebel et al., 2013; Zhang et al., 2014; Horowitz-Kraus et al., 2015; Borchers et al., 2019). However, other studies that have investigated the SLF, whether from a whole-brain or targeted approach, have failed to replicate this (Odegard et al., 2009; Welcome and Joanisse, 2014; Arrington et al., 2017), and a few studies have even reported negative FA-reading associations in the SLF (Carter et al., 2009; Frye et al., 2011).

The temporal sub-component of the SLF, the arcuate fasciculus (AF), has also been linked to reading performance (Rauschecker et al., 2009), and has often been analyzed separately from other SLF components due to its unique contributions to the language network (Catani et al., 2005). Similarly, FA reductions in dyslexia have been reported in the left AF (Klingberg et al., 2000; Deutsch et al., 2005; Vandermosten et al., 2012a; Marino et al., 2014; Christodoulou et al., 2017; Su et al., 2018), and its FA has been positively associated with reading skills (Klingberg et al., 2000; Deutsch et al., 2005; Yeatman et al., 2012a; Horowitz-Kraus et al., 2014; Christodoulou et al., 2017; Borchers et al., 2019). However, bilateral AF regions of higher FA in dyslexia have been identified (Žarić et al., 2018), and separate studies have reported negative FA-reading associations in the left AF (Yeatman et al., 2012a; Christodoulou et al., 2017; Huber et al., 2018).

Two fiber bundles that run under the SLF, the inferior longitudinal fasciculus (ILF) and inferior fronto-occipital fasciculus (IFO), serve to connect occipital and temporal-occipital areas to anterior temporal and frontal regions, respectively (Martino et al., 2010; Herbet et al., 2018), and have been identified as candidate reading tracts (Vandermosten et al., 2012a; Yeatman et al., 2013). The left ILF has exhibited increased FA in typically developing readers compared to dyslexic readers (Steinbrink et al., 2008; Marino et al., 2014; Su et al., 2018), and bilateral ILF FA has been positively related to reading performance (Steinbrink et al., 2008; Odegard et al., 2009; Feldman et al., 2012; Yeatman et al., 2012a; Lebel et al., 2013; Horowitz-Kraus et al., 2014; Zhang et al., 2014; Horowitz-Kraus et al., 2015). However, a few studies have found negative associations between FA and reading scores in the left ILF (Yeatman et al., 2012a; Huber et al., 2018), and one study has found increased left ILF FA in dyslexic individuals compared to their typically reading counterparts (Banfi et al., 2019). While studies have only identified positive correlations between FA and reading in bilateral IFO (Steinbrink et al., 2008; Odegard et al., 2009; Feldman et al., 2012; Lebel et al., 2013; Welcome and Joanisse, 2014; Zhang et al., 2014; Arrington et al., 2017), only a single study has found a reduction in left IFO FA in dyslexia (Steinbrink et al., 2008). It is worth noting that other investigations have also yielded null results in these tracts (Klingberg et al., 2000; Frye et al., 2011; Borchers et al., 2019).

The uncinate fasciculus (UF) is thought to contribute to the ventral orthographic pathways of reading, connecting temporal and orbitofrontal regions (Catani et al., 2002; Schlaggar and McCandliss, 2007). Despite reports of positive associations between FA and reading skills in bilateral UF (Odegard et al., 2009; Feldman et al., 2012; Welcome and Joanisse, 2014; Zhang et al., 2014; Arrington et al., 2017), the only significant group difference in FA that has been reported favored higher FA in dyslexia (Arrington et al., 2017). The same study also reported a negative correlation between FA and one of their reading measures in the right UF (Arrington et al., 2017). The splenium of the corpus callosum is also thought to contribute to reading, as it subserves interhemispheric communication between visual cortices (Putnam et al., 2010). Opposing results for both group (Frye et al., 2008; Marino et al., 2014) and continuous analyses (Frye et al., 2008; Odegard et al., 2009; Feldman et al., 2012; Lebel et al., 2013; Zhang et al., 2014; Huber et al., 2018) have been reported.

One theory of the etiology of dyslexia is the cerebellar hypothesis, which implicates cerebellar dysfunction in deficits of procedural learning and reading fluency (Nicolson et al., 2001; Nicolson and Fawcett, 2007; Stoodley and Stein, 2011). To this end, the contributions of the superior (SCP), inferior (ICP), and middle (MCP) cerebellar peduncles to reading have been investigated. The SCP contains efferent fibers that connect the deep cerebellum to inferior prefrontal regions involved in reading (Bruckert et al., 2020). The ICP and MCP contain primarily afferent fibers which connect the brainstem with the cerebellum. Such connections may facilitate automation of articulatory and oculomotor control (Bruckert et al., 2020). Bilateral SCP FA has exhibited negative relations to reading skill (Travis et al., 2015; Bruckert et al., 2020), while the left ICP has shown both positive (Borchers et al., 2019) and negative (Travis et al., 2015) associations. In the MCP, separate studies have paradoxically reported a higher FA in dyslexic readers compared to typically developing readers (Richards et al., 2008) as well as a positive association between FA and reading skills (Lebel et al., 2013; Travis et al., 2015).

Despite the general trend of higher FA relating to better reading and modest agreement in reading tract outcomes, a meta-analysis showed no evidence for systemic FA disruptions in dyslexia (Moreau et al., 2018). Small cohort sizes, inhomogeneous acquisition parameters, employment of different reading measures, variety of age groups, and diversity in processing and analytical methods may underlie the inconsistencies in past results (Moreau et al., 2018; Ramus et al., 2018; Schilling et al., 2021a). To address this, we leveraged a large database, the Healthy Brain Network (Alexander et al., 2017), to investigate white matter microstructural correlates of individual differences in single-word and single-nonword aptitude in children with diverse reading abilities. We additionally looked for tract-specific differences in FA between groups of children with and without reading disabilities. Considering the limited sensitivity of group difference analyses and the meta-analysis by (Moreau et al., 2018), we did not expect to find significant FA group differences. However, with the added specificity of using tract-based ROIs and sensitivity from an individual differences approach, we hypothesized that several tracts, particularly in the left hemisphere, would exhibit positive associations between FA and reading scores. Further, we performed exploratory analyses considering two possible factors that may have contributed to variable prior findings. First, we divided children into younger (ages 8 and below) and older (ages 9 and above) groups who are, respectively, learning to read versus reading to learn. Second, we examined whether findings differed between scores based on reading words, which includes memorized knowledge of specific words, versus reading pronounceable nonwords (pseudowords), which is a pure measure of knowledge of letter-sound correspondence.

## 2 Materials and Methods

### 2.1 Participants

We downloaded data from 1221 participants across the first 8 data releases of the Healthy Brain Network project (Alexander et al., 2017). Participants were all scanned at Rutgers University. All data were accessed in accordance with a data use agreement provided by the Child Mind Institute. The Healthy Brain Network project was approved by the Chesapeake Institutional Review Board (now called Advarra, Inc.; https://www.advarra.com/). The research team obtained written informed consent from participants ages 18 or older. For younger participants, written informed consent was collected from their legal guardians, while written assent was obtained from the participant. Full inclusion and exclusion criteria are described in the project’s publication (Alexander et al., 2017). Of note, each participant was fluent in English, had an IQ over 66, and did not have any physical or mental disorder precluding them from completing the full battery of scanning and behavioral examinations. Several behavioral and cognitive evaluations were collected as part of the project. Relevant to this study, participants completed the Test of Word Reading Efficiency 2^nd^ edition (TOWRE) (Torgesen et al., 1999) and the Edinburgh Handedness Inventory (EHI) (Oldfield, 1971).

The TOWRE consists of two subtests, Sight Word Efficiency (SWE) and Phonemic Decoding Efficiency (PDE). For Sight Word Efficiency, each participant is shown a list of words and asked to read the words aloud as quickly as possible. Raw scores are based on the number of words read correctly within the 45-second time limit and then converted to a standard score (population mean = 100, SD = 15). For Phonemic Decoding Efficiency, each participant is shown a list of pseudowords (pronounceable nonwords) and asked to read the pseudowords aloud as quickly as possible. Raw scores are based on the number of pseudowords read correctly within the 45-second time limit and then converted to a standard score (population mean = 100, SD = 15). The composite TOWRE score is the mean of the two standardized scores.

After quality control (QC), there were 686 participants ages 5-18 years old. We split these participants into two groups based on clinical diagnoses following the 5^th^ edition of the Diagnostic and Statistical Manual for Mental Disorders (Edition et al., 2013). There were 104 participants who were diagnosed with a “specific learning disability with impairment in reading” and assigned to the reading disability RD group, while the remaining 582 participants were assigned to the typically-reading (TR) group. It is worth noting that the specific criteria for diagnosing reading disabilities were not provided, and several participants with clinically low TOWRE scores - lower than 85 (Heath et al., 2006; Pugh et al., 2014; Johnston et al., 2016) - were not diagnosed with a reading disability (Figure 1).

**Figure 1:**
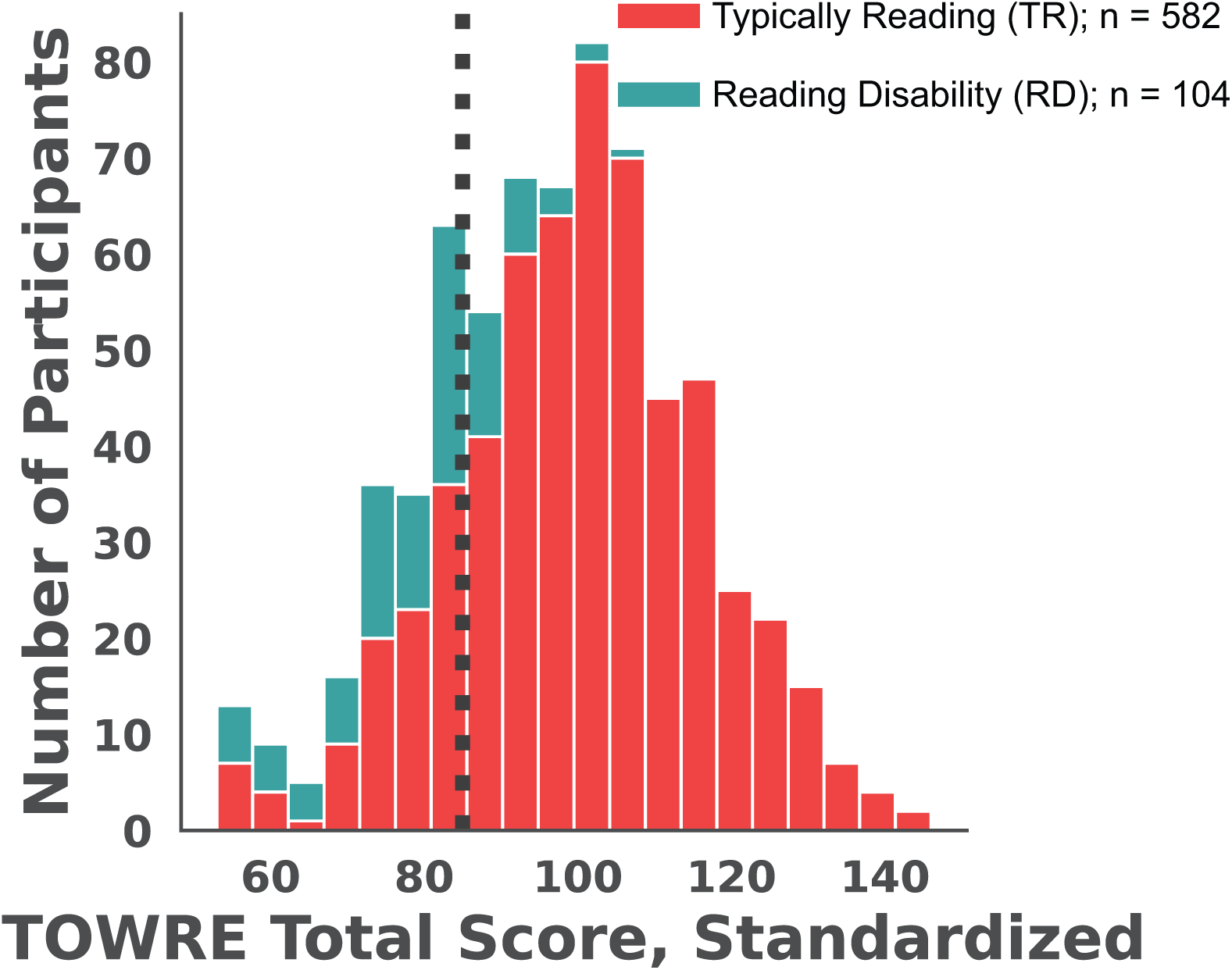
Distribution of standardized TOWRE composite scores across all participants. Bar colors denote whether the participant was formally diagnosed with a specific learning disability with impairment in reading (RD; teal) or not (TR; red). The black dotted line marks a TOWRE score of 85 which is conventionally used for diagnosing reading disabilities. TOWRE: Tests of Word Reading Efficiency.

We compared the ages, sex distribution, handedness, intracranial volume (ICV), and global FA (gFA) between the groups (Table 1). The TR group was slightly but significantly older than the RD group (p < 0.05; two-tailed Welch’s *t*-test). There were overall more males than females in the cohort, but the proportions of sexes did not significantly differ between groups (p > 0.05; *χ*^2^ test). Handedness did not differ between groups (p > 0.3; two-tailed Welch’s *t*-test). ICV in TR participants was greater than in RD participants (p < 0.05; two-tailed Welch’s *t*-test). The RD group had a greater global FA than the TR group (p < 0.05; two-tailed Welch’s *t*-test).

**Table 1:** Phenotypic information in the different reading proficiency groups. Values are listed as mean (standard deviation). For group comparisons, * denotes p < 0.05 (two-tailed Welch’s t-tests). EHI: Edinburgh Handedness Inventory, FA: fractional anisotropy, ICV: intracranial volume.

| **Metric** | **All** | **TR** | **RD** |
| --- | --- | --- | --- |
| *n* | 686 | 582 | 104 |
| Sex [M / F] | 447 / 239 | 387 / 195 | 60 / 44 |
| Age [years] | 10.69 (3.09) | 10.79 (3.18)* | 10.16 (2.52)* |
| Handedness [EHI] | 61.78 (50.69) | 60.92 (50.17) | 66.61 (53.74) |
| ICV [mm<sup>3</sup>] | 1500184 (160011) | 1505734 (160747)* | 1469124 (153688)* |
| Global FA | 0.38 (0.05) | 0.375 (0.052)* | 0.385 (0.043)* |

### 2.2 Neuroimaging Acquisition

Participants were scanned using a Siemens 3T Tim Trio scanner while wearing a standard Siemens 32-channel head coil A high resolution T1-weighted (T1w) sequence was collected with the following parameters: TR = 2500 ms, TE = 3.15 ms, Flip Angle = 8*°*, 0.8 mm isotropic voxel resolution. A diffusion kurtosis imaging scan was administered with the following parameters: TR = 3320 ms, TE = 100.2 ms, Flip Angle = 90◦, 1.8 mm isotropic voxel resolution, 1 b = 0 image, 64 noncollinear directions collected at b = 1000 s/mm^2^ and b = 2000 s/mm^2^. A pair of PEpolar fieldmaps were collected before the diffusion scan to quantify magnetic field inhomogeneity. Detailed scanner protocols are published on the Healthy Brain Network project website (http://fcon_1000.projects.nitrc.org/indi/cmi_healthy_brain_network/File/mri/HBN_RU_Protocol.pdf).

### 2.3 Neuroimaging Preprocessing

Results included in this manuscript come from preprocessing performed using *QSIPrep* 0.13.0RC1 (Cieslak et al., 2021, RRID:SCR_016216) (https://qsiprep.readthedocs.io/en/latest/) which is based on *Nipype* 1.6.0 (Gorgolewski et al., 2011, 2018, RRID:SCR_002502). Many internal operations of *QSIPrep* use *Nilearn* 0.7.0 (Abraham et al., 2014, RRID:SCR_001362) and *Dipy* 1.3.0 (Garyfallidis et al., 2014, RRID:SCR_000029). Much of the text in the following two sections was provided by *QSIPrep* under a CC0 license so it may be included in a manuscript for the sake of transparency and reproducibility. We made minor changes for succinctness.

#### 2.3.1 Anatomical Preprocessing

The T1w image was corrected for intensity non-uniformity (INU) with N4BiasFieldCorrection (Tustison et al., 2010), distributed with *ANTs* 2.3.3 (Avants et al., 2008, RRID:SCR_004757), and used as the T1w-reference throughout the workflows. The T1w-reference was then skull-stripped with a *Nipype* implementation of the antsBrainExtraction.sh workflow (from *ANTs*), using OASIS30ANTs as the target template. Brain tissue segmentation of cerebrospinal fluid (CSF), white matter (WM) and gray-matter (GM) was performed on the brain-extracted T1w using fast (Zhang et al., 2001, *FSL* 5.0.9, RRID:SCR_002823). Additionally, brain surfaces were reconstructed using recon-all (*FreeSurfer* 6.0.1) (Dale et al., 1999, RRID:SCR_001847).

#### 2.3.2 Diffusion MRI Preprocessing

MP-PCA denoising as implemented in *MRtrix3*’s dwidenoise (Veraart et al., 2016) was applied with a 5-voxel window. After MP-PCA, Gibbs unringing was performed using *MRtrix3*’s mrdegibbs (Kellner et al., 2016). Following unringing, B1 field inhomogeneity was corrected using dwibiascorrect from *MRtrix3* with the N4 algorithm (Tustison et al., 2010). After B1 bias correction, the mean intensity of the diffusion-weighted imaging (DWI) series was adjusted so all the mean intensity of the b = 0 images matched across each separate DWI scanning sequence.

*FSL*’s (version 6.0.3:b862cdd5) eddy function was used for head motion correction and Eddy current correction (Andersson and Sotiropoulos, 2016). The function was configured with a *q*-space smoothing factor of 10, a total of 5 iterations, and 1000 voxels used to estimate hyperparameters. A linear first level model and a linear second level model were used to characterize Eddy current-related spatial distortion. *q*-space coordinates were forcefully assigned to shells. Field offset was attempted to be separated from participant movement. Shells were aligned post-eddy. eddy’s outlier replacement was run (Andersson et al., 2016). Data were grouped by slice, only including values from slices determined to contain at least 250 intracerebral voxels. Groups deviating by more than 4 standard deviations from the prediction had their data replaced with imputed values. Data were collected with reversed phase-encoded blips, resulting in pairs of images with distortions going in opposite directions. Here, b = 0 reference images with reversed phase encoding directions were used along with an equal number of b = 0 images extracted from the DWI scans. From these pairs the susceptibility-induced off-resonance field was estimated using a method similar to that described in (Andersson et al., 2003). These susceptibility maps were ultimately incorporated into the Eddy current and head motion correction interpolation. Final interpolation was performed using the jac method. Slicewise cross correlation was calculated. The DWI time-series were resampled to ACPC with 1.2mm isotropic voxels.

### 2.4 Tract Segmentation

We used *TractSeg* version 2.3 (Wasserthal et al., 2018, 2019) (https://github.com/MIC-DKFZ/TractSeg), a deep-learning based white matter segmentation method, to reconstruct fiber bundles. We chose this method due to its favorable balance between the accuracy of manual fiber tracking and objectivity of atlas-based methods (Genc et al., 2020). This involved the following steps: First, we reoriented images (preprocessed T1w, DWI, and brain mask) to the *FSL* standard space with `fslreorient2std` and accordingly corrected the diffusion gradient table with *MRtrix3*’s `dwigradcheck`. We used the *MRtrix3* commands `dwi2tensor` and `tensor2metric` to fit the diffusion tensor with an iterative weighted least-squares algorithm (Basser et al., 1994; Veraart et al., 2013) and produce the FA map. Multi-tissue fiber response functions were estimated using *MRtrix3*’s dhollander algorithm (Dhollander et al., 2016, 2019). Fiber orientation densities (FODs) were estimated via multi-shell multi-tissue constrained spherical deconvolution (CSD) (Tournier et al., 2004, 2008; Jeurissen et al., 2014). FODs were intensity-normalized using mtnormalize (Raffelt et al., 2017). The first three principal FOD peaks were extracted and used as inputs into *TractSeg*’s convolutional neural network. We produced tract segmentations for the following 9 bilateral tracts: AF, SLF (I, II, and III), ILF, IFO, UF, SCP, and ICP. We also segmented the MCP and splenium of the corpus callosum (CC 7 by *TractSeg* naming convention), leading to a total of 20 fiber bundles to analyze. We extracted the average FA in each tract by calculating the average intensity of the intersection between the tract’s segmentation mask with the participant’s FA map. As a *post-hoc* analysis, we analyzed FA at several points along the fiber bundle (Yeatman et al., 2012b) instead of using a tract average. For this procedure, streamlines were generated for each tract. This method and its results are described in the Supplementary Materials.

### 2.5 Statistical Analysis

Several covariates were included in the public data set as phenotypic information, including sex (binary 0 or 1), age (in years), and handedness (EHI score; from -100 to 100). We additionally extracted estimated total intracranial volume (ICV) from *FreeSurfer* reconstructions, as well as the global white matter FA (gFA). We calculated gFA by binarizing the white matter probabilistic segmentation from *QSIPrep’s* anatomical workflow at a threshold of 0.5, and averaging the FA intensity within the resulting white matter mask. We did not define gFA on a whole-brain scale as that would have introduced variance from potential differences in white matter volumetric proportions. Given the novelty of our analytical tools and recency of the data set, we first ran Spearman correlations between these phenotypic and neuroimaging-derived metrics, probing well-established relationships in our data as a way of validating our approach and informing our choice of model parameters (Figure 2).

**Figure 2:**
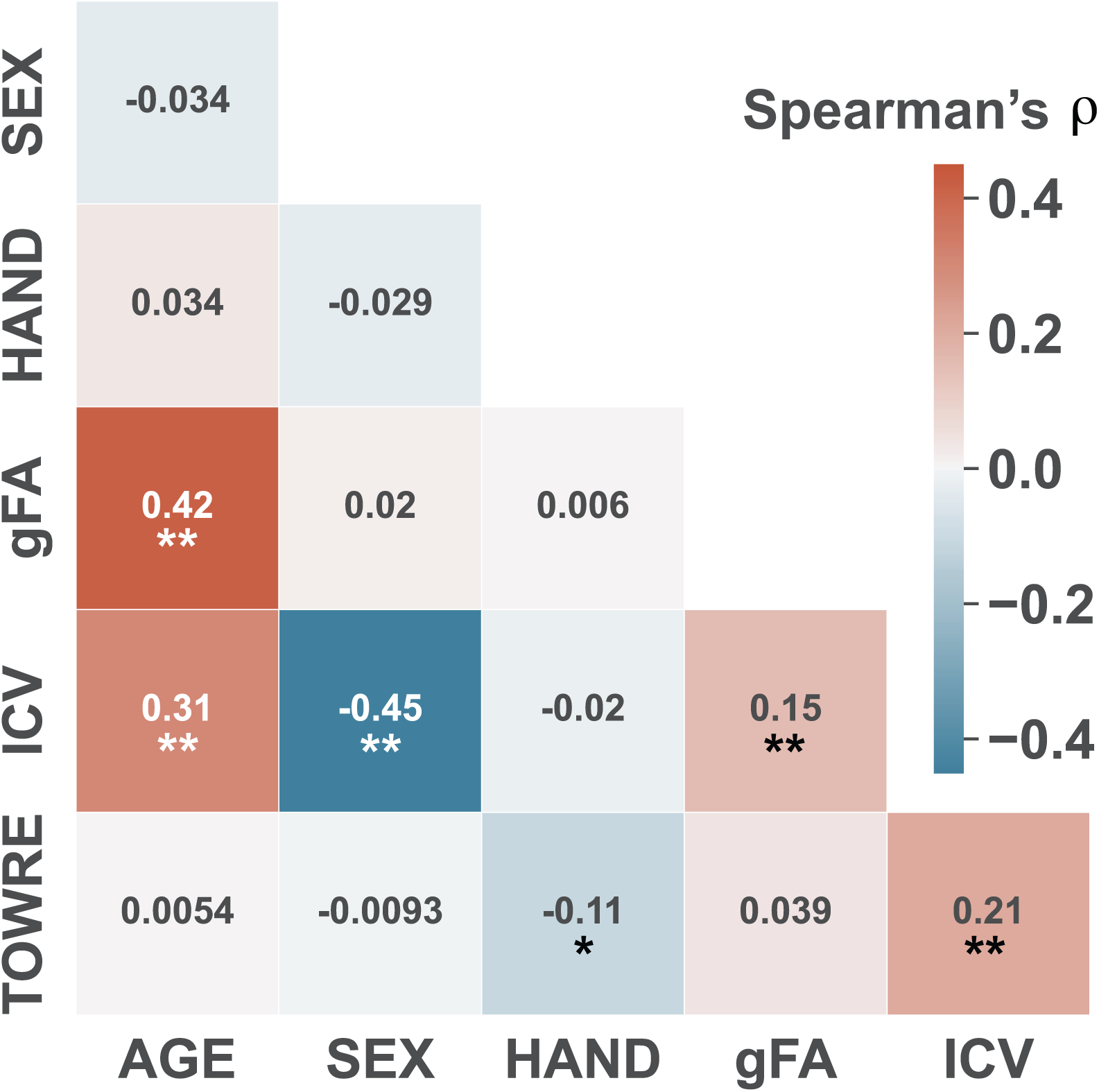
Correlations between phenotypic and neuroimaging-derived metrics. The color of each cell corresponds to the Spearman’s *ρ* of the correlation between the metrics in the respective row and column. * denotes p < 0.01, ** denotes p < 0.001, adjusting for multiple comparisons (Benjamini-Hochberg FDR). gFA: global fractional anisotropy, HAND: Edinburgh Handedness Inventory score, ICV: intracranial volume, TOWRE: Tests of Word Reading Efficiency (age-standardized composite score).

For each tract, we computed both a correlation between mean FA and composite TOWRE scores (skipped correlation (Wilcox, 2004; Rousselet and Pernet, 2012) with Spearman’s *ρ*) and group difference (Welch’s *t*-test) between mean FA values in the RD and TR groups. Before running statistical tests, we removed any participant whose tract FA was less than 0.2, as this indicated either a tract segmentation containing significant amount of non-white matter or a tract being out of the field-of-view (this happened most often for the UF in orbitofrontal cortex and the ICP in the cerebellum). This accounted for no more than 4 participants in any given tract. For group difference analyses, we linearly regressed out the effects of sex, handedness, age, global FA, and the interaction between group and sex on mean tract FA. For correlation analyses, we regressed out sex, handedness, and ICV. Standardization of TOWRE scores accounted for differences in age. Covariates were demeaned and rescaled to unit variance before regression. 40 statistical inferences were made (20 tracts × 2 measures per tract). We used Benjamini-Hochberg FDR correction to account for multiple comparisons (Benjamini and Hochberg, 1995). Statistical tests were executed by the Python package Pingouin (0.5.0) (Vallat, 2018) and visualized with the Seaborn package (0.11.2) (Waskom, 2021).

We conducted several *post-hoc* exploratory analyses to probe additional factors that may contribute to variation in prior DTI studies of reading. Recognizing that reading processes may differ between early and late-stage readers, we divide the cohort into two age brackets based on a threshold of 9 years old. This resulted in 455 older participants and 231 younger participants. Additionally, to account for differences in neurocognitive mechanisms between single-word and single-nonword reading, we also ran correlations against the SWE and PDE sub-scores. We used Benjamini-Hochberg FDR correction within each of these secondary analyses to account for multiple testing. Finally, we ran analyses of tract profiles (see Supplementary Materials), in which inferences are calculated on small segments along the length of a bundle.

### 2.6 Data Inclusion and Quality Control

Of the original 1221 participants, 862 participants had all the necessary neuroimaging data (T1w, diffusion, and fieldmap) and were able to be run through *QSIPrep* and *TractSeg* without errors. Three of those participants were excluded for having ubiquitously sparse fiber bundle reconstructions. Two additional participants had misaligned anatomical outputs from *QSIPrep*, and were excluded because gFA could not be calculated for them. Additionally, 152 of the remaining participants were missing either *FreeSurfer* reconstructions or necessary phenotypic data. An additional 16 participants were excluded for being over 18 years old. For the remaining participants, due to the volume of images included in the cohort, visual inspection of each DWI image was not practical. We instead adopted an automated QC procedure described in (Yeh et al., 2019), which has been integrated into the outputs of *QSIPrep*. This involves rejecting a scan if it had different or incomplete scanning acquisitions (no participants excluded) or if over 0.1% of slices (9 slices at 72 slices/volume × 128 diffusion volumes) had significant signal dropout based on changes in slice-wise correlation (3 participants excluded). Therefore, a total of *n* = 686 participants were analyzed.

### 2.7 Data and Code Availability

Neuroimaging and phenotypic data can be collected following directions on the Healthy Brain Network data portal (http://fcon_1000.projects.nitrc.org/indi/cmi_healthy_brain_network/index.html) after signing a data use agreement. We cannot distribute this data publicly. All code and instructions for preprocessing neuroimaging data and running the statistical models can be found at https://github.com/smeisler/Meisler_ReadingFA_Associations. With minimal modification, the preprocessing code should be able to run on most BIDS-compliant data sets using the SLURM job scheduler (Yoo et al., 2003). Some softwares we used were distributed as Docker (Merkel, 2014) containers, then compiled and run with Singularity (3.6.3) (Kurtzer et al., 2017):

- *QSIPrep* 0.13.0RC1 (`singularity build qsiprep.simg docker://pennbbl/qsiprep:0.13.0RC1`)
- *TractSeg* 2.3 (`singularity build tractseg.simg docker://wasserth/tractseg:master`)
- *MRtrix* 3.0.3 (`singularity build mrtrix.simg docker://mrtrix3/mrtrix3:3.0.3`)
- *FSL* 6.0.4 (`singularity build fsl.simg docker://brainlife/fsl:6.0.4-patched`)

We encourage anyone to use the latest stable releases of these softwares.

## 3 Results

### 3.1 Correlations Between Phenotypic and Neuroimaging Measures

We examined relationships between phenotypic and neuroimaging metrics (Figure 2). There was a significant and positive association between ICV and composite TOWRE scores (Spearman’s *ρ* = 0.21, p < 0.001). There was also a significant and positive association between ICV and gFA (Spearman’s *ρ* = 0.51, p < 0.001). In relation to age (development), there were strong positive correlations between age and gFA (Spearman’s *ρ* = 0.42, p < 10*^−^*^30^) and between age and ICV (Spearman’s *ρ* = 0.31, p < 0.001). A negative correlation between sex (M = 0, F = 1) and ICV suggests that females had, on average, smaller brain volume than males in this cohort (Spearman’s *ρ* = -0.45, p < 10*^−^*^30^). Interestingly, we found a modest negative correlation between handedness quotients and TOWRE scores (Spearman’s *ρ* = -0.11, p < 0.01), suggesting that right-handedness was related to worse reading ability. However, this comparison was unbalanced due to the high preponderance of right-handed participants (Table 1). SWE and PDE scores were highly correlated (Spearman’s *ρ* = 0.79, p < 10*^−^*^30^).

### 3.2 Mean FA-TOWRE Associations and Group Differences

Among all participants, no tract exhibited either significant associations between FA and TOWRE composite scores or significant FA differences between RD and TR groups. Correlation coefficients approximately ranged from -0.036 to 0.036, and t-statistics ranged from -0.92 to 0.24 (with a positive statistic indicating TR > RD). No significant correlations or group differences were observed when restricting analyses to specific age brackets. However, associations tended to be stronger and more positive, albeit still modest, among the older readers, with the right SLF I exhibiting a correlation coefficient of 0.09, corresponding to a p-value of 0.07.

### 3.3 TOWRE Sub-Score FA Correlations

No correlations between SWE scores and mean tract FA were significant within either age bracket. Significant associations between PDE scores and FA were significant only among children ages 9 years or older (Figure 3). In this age bracket, FA in the right SLF I (Spearman’s *ρ* = 0.10, p = 0.038) and left ICP (Spearman’s *ρ* = 0.09, p = 0.048) were positively associated with PDE scores. Neither of the results remained significant after multiple comparison correction.

**Figure 3:**
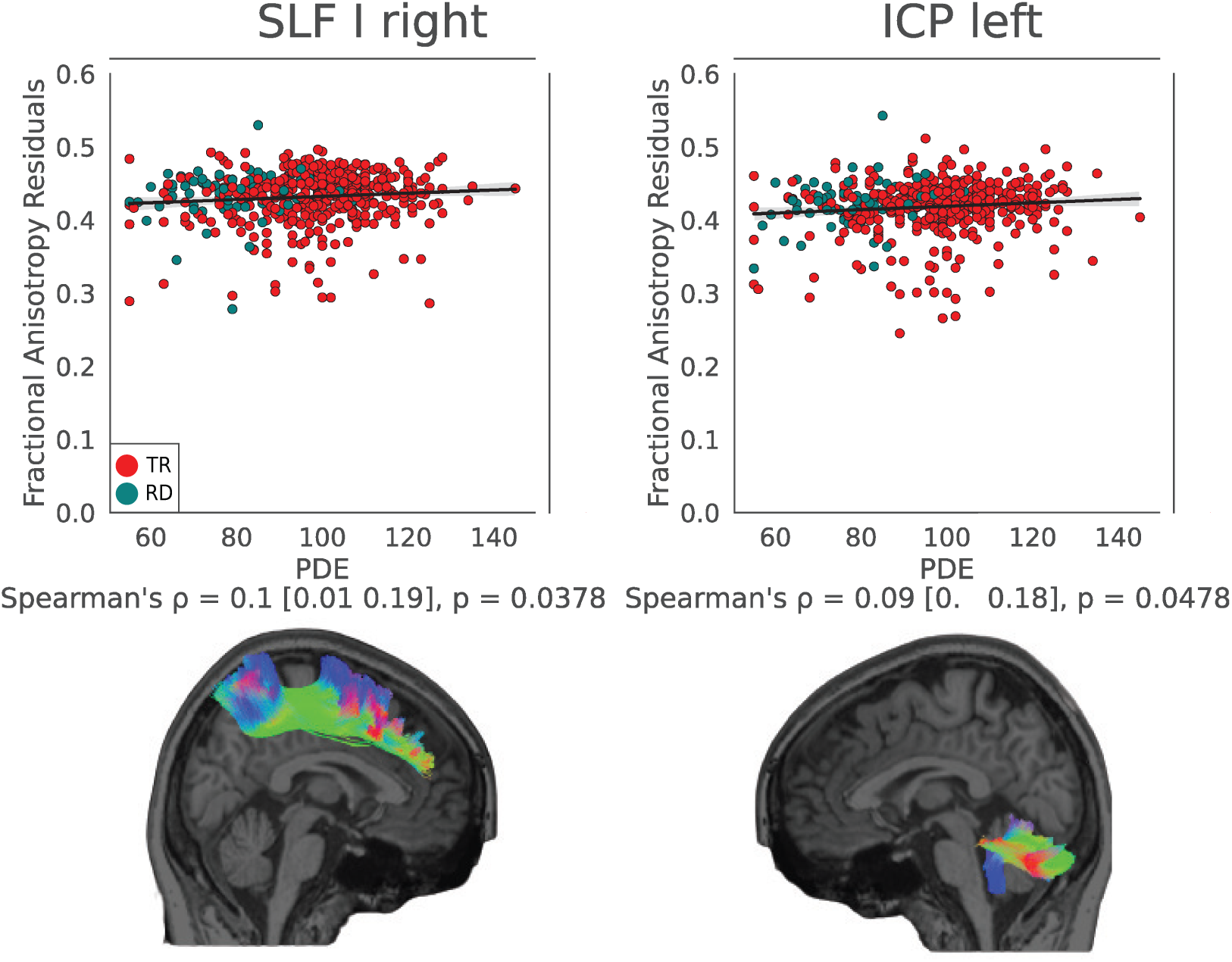
Significant correlations between mean tract FA and TOWRE Phonemic Decoding Efficiency (PDE) scores among older participants (*n* = 455). The dots map participants’ age-standardized PDE scores against average tract FA after regressing out sex, handedness, and intracranial volume. Teal denotes a RD participant, and red denotes TR participants. The black line represents the best fit line across all participants, and the shaded region surrounding it depicts the 95% confidence interval of the fit. Summaries of the statistical tests are written out below each plot, including test coefficients, 95% confidence intervals surrounding the test coefficient, and uncorrected *p*-values. Tract shapes and anatomical image come from a single subject. Tract color relates to streamline direction. SLF: superior longitudinal fasciculus, ICP: inferior cerebellar peduncle, TR: Typically reading group, RD: reading disability group.

## 4 Discussion

This study is the first large-scale investigation of tract segmentation-derived white matter microstructural associations with single-word and single-nonword reading abilities across a diverse pediatric data set. The size of our participant cohort was large relative to other DWI studies, particularly those relating to reading aptitude. We used high-quality publicly available data and state-of-the-art analytical methods, showcasing the rapid advances in diffusion imaging acquisition and processing techniques. No tracts exhibited group differences in average FA between children with and without reading disabilities, even when considering more specific age brackets. Positive correlations between FA and reading abilities were only significant in the right SLF and left ICP when considering older participants and PDE scores, and even then, effect sizes were modest and did not survive multiple comparison correction.

Our hypotheses were supported in that, overall, there were no significant group differences in tract-specific FA values between children with versus without reading disabilities or and no significant correlations between reading scores and tract-specific FA values.

The present findings are surprising in several ways, especially given the exceptional statistical power of the present sample. Firstly, we found that children with reading disability had higher global FA values than their typically reading counterparts. This differs from the frequent focus in dyslexia research on reduced FA in white matter tracts connecting the major occipito-temporal, temporal-parietal, and frontal nodes of the reading network. Secondly, while the the use of age-standardized reading scores controlled for simple age-related gains in reading, but there was little relation between composite TOWRE scores and white matter pathway FA. Using G*Power 3.1 (Faul et al., 2009), we conducted a power analysis to see how many participants would be needed for the strongest correlation achieved in the primary analysis to be significant. At that effect size (0.036), with a desired *α* = 0.05 and power of 0.8, 4766 participants would be required. Even if one could recruit that many participants, such a low effect size presents challenges in interpretation. Significant effect sizes were only found in our *post-hoc* exploratory analyses, and even these were modest. These secondary analyses revealed significant correlations only when using PDE scores, despite PDE and SWE scores reading being highly related, and only when examining older children (ages 9-18). At the very least, the significant correlations between FA and PDE scores were positive, which is consistent with most prior neuroanatomical studies of reading.

Given the left-lateralized dominance of reading and language brain circuitry, we did not expect FA associations to be present in the right hemisphere, at least when considering a cohort that is predominantly composed of typical readers. There has been some evidence that poor readers utilize right-hemisphere pathways to compensate for limitations of the left-hemisphere pathways that are typically most associated with single-word reading. Several studies have reported enhanced right-hemisphere brain activation in individuals with dyslexia while reading, which has been interpreted as a compensatory mechanism (Milne et al., 2002; Temple et al., 2003; Hoeft et al., 2011; Waldie et al., 2013). Greater FA in right-hemisphere white matter pathways, in particular the right SLF, has been associated with better reading outcomes in children with dyslexia (Hoeft et al., 2011) and in young pre-reading children with low scores on assessments that predict future reading difficulty (Zuk et al., 2021). Positive correlations between right SLF FA and reading abilities have also been observed among typical readers (Horowitz-Kraus et al., 2015). However, opposing results have been reported for correlations between right SLF FA and pseudoword reading ability in particular (Steinbrink et al., 2008; Frye et al., 2011).

The TOWRE composite score was the average between real-word (SWE) and nonword (PDE) reading ability. These two measures are typically correlated, and both are impaired in dyslexia, but there are distinctions between the two measures. Single-word reading partially reflects reading experience with real words, including memorization of words. Single nonword reading, although less directly related to reading experience, provides a more pure measure of letter-sound decoding of print. We found significant, albeit small, positive correlations between FA and nonword reading scores only. This finding suggests that white matter microstructure may be especially salient for letter-sound decoding of print.

We found that age was related to finding associations between FA and nonword reading skill such that only in older children (ages 9 and older) did some tracts show a positive correlation between PDE scores and FA. Such brain-behavior relations might differ in younger children who are learning to read single words and older children who have mastered single-word reading skills and are more focused on reading comprehension of advanced texts. Indeed, there is evidence that learning a skill leads to an initial increase in FA in putative white matter regions followed by a decline when reaching proficiency (Scholz et al., 2009), and similar early changes in white matter have been reported in mathematical abilities in dyslexia (Koerte et al., 2016), musicianship (Schmithorst and Wilke, 2002), and dance (Hänggi et al., 2010).

We, however, observed the opposite relation such that higher white matter FA was associated with nonword reading skill only in older children with more advanced reading skills. It is possible that we did not observe any significant relationships in younger readers due to reduced statistical power. In addition, our results may suggest that variation of microstructural properties of white matter does not relate to the etiology of reading ability, but rather to the long-term consequence of one’s reading ability. That is, in older participants, lower FA among poor readers could be the result of having lived for years with reading difficulties (Protopapas and Parrila, 2018, 2019). A similar association of gray matter volume and reading skills was observed only in older participants in a different large-scale study (Torre and Eden, 2019). In that study, reading ability was primarily correlated with right superior temporal gyrus volume among males. Alternatively, differential reading instruction in early grades may confound reading experience and reading skill in the younger children with reading disability.

There is debate surrounding whether dyslexia can properly be described as a neurodevelopmental disorder. Population reading skills tend to follow a normal distribution, even when including those with developmental dyslexia (Figure 1). One may argue that dyslexia is not a neurodevelopmental disorder, but rather a way to group those on the lower end of the reading skills distribution (or bell curve). From this perspective, both reading differences and associated brain differences should lie along a continuum (Protopapas and Parrila, 2018, 2019).

At the simplest interpretation of this argument, being that the behavioral and neuroimaging indices lie on the same continuum, findings that would most strongly align with this view would be if reading-FA correlations were positively correlated across the entire participant cohort. We found no such association, either globally or within tracts. This suggests that the continua relating reading aptitude and FA are more complex than a shared linear relationship. Indeed, FA-behavior relationships may not be generalizable across domains and may index various neural and behavioral measures (Lazari et al., 2021).

This is the first study of reading skills that has used *TractSeg*, which has been shown to outperform several white matter segmentation methods (Wasserthal et al., 2018; Schilling et al., 2021b). Newer tract-based approaches, as employed here, represent a paradigm shift from several earlier DTI studies. Many previous papers of FA-reading relationships, particularly before the publication of Vandermosten and colleagues’ review of DTI applications to reading (Vandermosten et al., 2012b), performed whole-brain voxel-based analyses (VBA). VBA sensitivity suffers from stricter multiple-comparison correction across the entire brain. In addition, VBA methods tend to be less precise due to their being performed on a group-averaged image because the shape of long-range fiber bundles varies among people (Yeatman et al., 2011; Wassermann et al., 2011). Spatial smoothing and affine transformation to MNI before group analysis in VBA may obfuscate unique properties of a participant’s anatomy (Christensen et al., 1997). Significant findings from VBA are not always assigned to tracts. In fact, a single voxel may contain multiple fiber bundles, so it is possible early studies may have incorrectly ascribed significant voxels to fiber bundles due to not also considering the primary diffusion directionality in these areas. Tract-Based Spatial Statistics (TBSS) (Smith et al., 2006), a method that improves upon traditional VBA by restraining analyses to a skeletonized white matter voxel map, also suffers in tract localization (Tsang et al., 2010). Methodological examinations of TBSS have revealed bias in the FA map used for spatial normalization, sensitivity to pathologies and noise that affect brain anatomy, incorrect voxel-to-tract assignments (Bach et al., 2014), and instability dependent on tensor-fitting methods (Maximov et al., 2015). VBA can still be valuable as a data exploratory tool, but tract-based methods likely provide for superior sensitivity and reduced ambiguity of fiber bundle localization (Ramus et al., 2018). These benefits have already been observed in multiple clinical populations (Kamagata et al., 2013; Kuchling et al., 2018; Wasserthal et al., 2020; Forkel et al., 2021).

With these limitations in mind, it may not be surprising that tract-based methods have yielded some opposing findings to voxel-based methods. For example, in the left AF, tract-based studies have yielded observations of lower FA being associated with better reading scores (Yeatman et al., 2012a; Christodoulou et al., 2017; Huber et al., 2018), as well as findings of higher FA in dyslexic readers compared to typical readers (Žarić et al., 2018). However, tractography applications to reading are still in their infancy, and different tractography measures may provide different insights. For example, CSD methods are better at resolving fiber bundles that pure DTI methods have difficulty with, such as the right AF (Catani et al., 2007; Yeatman et al., 2011; Zhao et al., 2016). The field may best be served by harmonizing diffusion acquisition and analysis protocols (Ramus et al., 2018; Cieslak et al., 2021; Schilling et al., 2021a,b), particularly scanning with multiple shells including high b-values (at least b = 1300) to optimally use CSD.

While our largely null results stand in opposition to previous studies reported significant FA-reading associations or group differences, a recent meta-analysis of VBA studies has suggested that these findings are not robust (Moreau et al., 2018). The present study extends this in a tract-based paradigm, using a single data set to mitigate concerns of variability from scanning protocol confounding results (Schilling et al., 2021b). Our findings also underscore the importance including global confounds to isolate local from global effects (Ramus et al., 2018). Intracranial volume and TOWRE scores were significantly related (Figure 2), and groups differed in global fractional anisotropy (Table 1). If we had not included these factors in our models, we would have likely reported significant effects in several bilateral tracts, despite the effects being driven by global differences.

Unrelated to reading skill, we found strong correlations between age and global FA. Indeed, the present study is the largest investigation of linear white matter development patterns to date (for a review, see (Lebel et al., 2019); for a similarly-sized group-based analysis, see (Chiang et al., 2011)). Our findings are consistent with prior reports of linear FA-age relationships during child development (Mukherjee et al., 2002; Schmithorst et al., 2002; Barnea-Goraly et al., 2005; Bonekamp et al., 2007; Muetzel et al., 2008), although more recent studies suggest an exponential trajectory (Lebel et al., 2008; Tamnes et al., 2010; Taki et al., 2013; Simmonds et al., 2014) which we did not evaluate in this study. Several of our other findings unrelated to reading skill are also consistent with well-established trends, including worse readers having smaller brain volumes (Ramus et al., 2018), males having larger brain volumes than females (Courchesne et al., 2000), and brain volumes correlating positively with global FA (Takao et al., 2014). These agreements with prior studies support the quality of the DWI acquisition and the validity of our analytic method.

The results of this study should be interpreted in the context of some limitations. Firstly, the composition of the participant cohort is unique in that most participants in the Healthy Brain Network have at least one psychological, neurodevelopmental, or learning disorder (Alexander et al., 2017). Incentives for participating in the study included a cash reward as well as a free behavioral examination, and advertisements were targeted towards parents who may have been concerned about their child’s psychological state. RD participants comprised approximately 15% of the cohort. This figure exceeds estimates of dyslexia prevalence which tend to range from 5-10% (Roongpraiwan et al., 2002). Secondly, we used FA as our tract metric of interest since it has largely been interpreted as a measure of structural integrity. There is often an implicit assumption that a higher FA may relate to more myelination, which in turn may increase synaptic efficiency (Lebel and Deoni, 2018). While myelination certainly modulates FA, other factors such as axonal diameter, density, and coherence also influence it (Beaulieu, 2009; Shemesh, 2018; Friedrich et al., 2020). The dueling mechanisms underlying FA may confound results, as both axonal pruning and increased myelination may relate to typical brain development but have opposing effects on FA (Yeatman et al., 2012a). FA is also prone to underestimation and noise in voxels that contain both gray and white matter or crossing fibers (Oouchi et al., 2007), which would especially confound measurements of the corpus callosum. Analysis of T1 relaxation time in tracts using quantitative MRI may better quantify the degree of myelination per se (Lutti et al., 2014; Schurr et al., 2018). Thirdly, we could not account for FA alterations related to nonverbal IQ, since this information, in the form of the Kaufman Brief Intelligence Test (Kaufman, 1990), Second Edition, was not available for most participants. Next, our significant results should be interpreted with some caution, as they originated from exploratory analyses for which we did not develop *a priori* hypotheses. Finally, while our results lend themselves to theories of the etiology of dyslexia, we cannot interpret our findings to make causal statements of white matter microstructural properties’ connection to reading disabilities. Longitudinal interventional studies should continue to be performed to better understand this relationship (Huber et al., 2018). Multivariate pattern analysis could shed insight into the relative contributions of different tract microstructural properties to classifying reading aptitude, similar to what was done in the study by (Cui et al., 2016).

The tract-profile approach employed in the Supplementary Materials, in which FA is sampled at several points across the length of the tract, allows researchers to detect group differences and correlations that may be localized to certain portions of a tract, even if such tests using the mean FA may be null (Yeatman et al., 2012a). Using this method, we found segments of greater FA in the TR group within the left SLF and UF (Figure S1). However, it is unclear if these localized phenomena are a result of true structural differences in a given tract, or more global structural differences such as inhomogeneities in the location of a crossing fiber (Ramus et al., 2018). It also is not well-established what effect, if any, a local disruption in FA might have on synaptic efficiency between disparate areas on a fiber bundle. This presents a trade-off between anatomical specificity and interpretability. For these reasons, we keep the tract profile analyses in the Supplementary Materials.

## 5 Conclusion

In several ways, this study of the relation between white matter microstructure and single-word reading ability had features that ought to have yielded conclusive findings, but the study raises some open questions. The number of participants was exceptionally large, the TOWRE is a widely used and well-established measure of single-word reading ability, and the analyses took advantage of progress in DWI analytic methods. Our largely null findings, while not consistent with several individual DTI studies of reading, are consistent with recent larger-scale meta-analyses of neuroanatomical studies of dyslexia (Moreau et al., 2018; Ramus et al., 2018). Without replicating individual processing pipelines, it is difficult to identify precise reasons previous results were or were not consistent with this study. In this respect, we believe that large-scale population studies and longitudinal studies with reproducible methods should be emphasized to account for inter-participant variability not reliably represented in small cross-sectional samples. Only future research can resolve the apparent contradictions among findings, but one possibility is that there is greater diversity among readers than has been heretofore imagined, perhaps combined with developmental variation from infancy to beginning readers to more advanced readers.

## CRediT Authorship Contributions

**Steven L. Meisler:** Conceptualization, Formal analysis, Investigation, Methodology, Software, Visualization, Data Curation, Writing - original draft, Writing - review & editing. **John D.E. Gabrieli:** Conceptualization, Supervision, Writing - original draft, Writing - review & editing.

## Acknowledgements

We thank the Healthy Brain Network team for their diligence in collecting the data and generosity in sharing it. We thank all of the participants and their families for volunteering their time to participate in the study. We thank Evelina Fedorenko, Nadine Gaab, and Tyler Perrachione for their constructive criticism of the manuscript. Finally, we thank Amanda O’Brien for proofreading the manuscript.

## Funding

This work was funded by the National Institutes of Health (grant numbers 5T32DC000038-29 and 5T32DC000038-30), the Halis Family Foundation, and Reach Every Reader, a grant supported by the Chan Zuckerberg Foundation.

## Conflicts of Interest

Declarations of interest: none

## Notes

https://github.com/smeisler/Meisler_ReadingFA_Associations

### Competing Interest Statement

The authors have declared no competing interest.

### Summary of Updates

We included a global confound (intracranial volume) for our analyses, and introduced a group analysis of FA in children with and without reading disability.

