## Supplementary Materials for "A Large-Scale Investigation of White Matter Microstructural Associations with Reading Ability"

### Tract Profile Analysis

Recently, tract profiles have been proposed as a more sensitive method to analyze microstructural properties of a fiber bundle (Yeatman et al., 2012). For this analysis, FA values were mapped to segments of reconstructed tract streamlines. To make the streamlines for each tract, we fed the FOD peaks into *TractSeg* to create binary masks of the tract, the ROIs that define its endpoint, and a corresponding tracking output map (TOM) (Wasserthal et al., 2018). These were used as inputs to *TractSeg*'s pre-trained convolutional neural network (Wasserthal et al., 2019, 2020). We generated a fixed large number (5000) of fibers per instance to account for the stochastic nature of reconstruction. This reduces inter-run variability. Using an approach based on *Dipy*'s Bundle Analytics framework (Chandio et al., 2020), FA values were calculated for 100 nodes along the length of the tract. At each node, we used pre-compiled *TractSeg* code to employ a non-parametric permutation-based statistical comparison (Nichols and Holmes, 2002) with 5000 iterations to look for either correlations (Pearson's $r$) between node FA and standardized TOWRE composite scores, or group differences (two-sample $t$-test) between group FA values. This permutation approach accounts for multiple comparisons *within* a tract given the correlative structure between adjacent nodes (Yeatman et al., 2012). We additionally imposed FWE correction for multiple comparisons *between* the 20 tracts, after correcting for within-tract comparisons. Covariate choices for group and correlation tests were the same as in the tract-averaged analyses. No correlations between node FA and composite TOWRE scores were significant after within-tract multiple comparison correction. Small regions of TR > RD were present in the left SLF and UF after correcting for multiple comparisons within-tract. These did not survive between-tract multiple hypothesis correction, however (Figure S1).

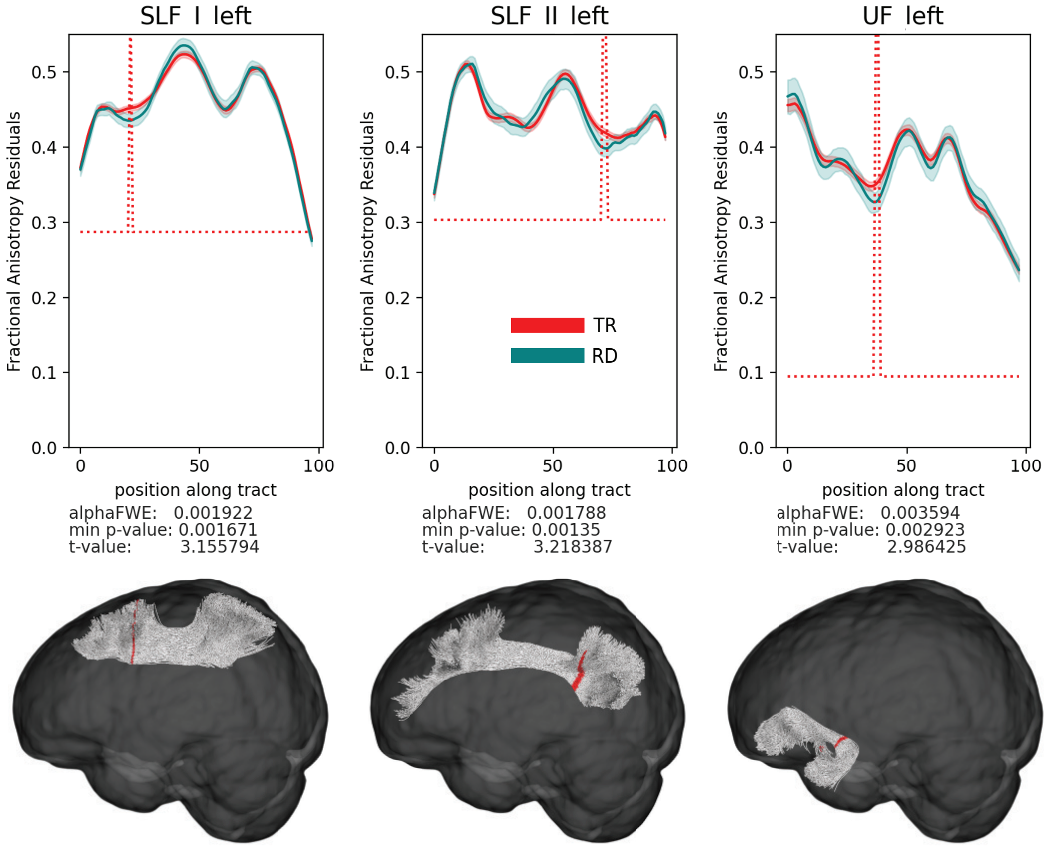


Figure S1: Significant tract-profile group comparison analyses. FA residuals for the TR and RD group are plotted as a function of position along the fiber bundle, after regressing out sex, age, handedness, global fractional anisotropy, and sex-group interactions. Shaded regions surrounding the curves denote 90% confidence intervals. The red dotted line extends above the curves at any segment where the two-sample t-test is significant after FWE correction within the tract Below each plot, the FWE-corrected alpha threshold (correcting for the 100 analyses within each tract), highest test coefficient achieved, and its associated p-value are listed. Accompanying each plot, tracts are highlighted red at significant segments. Tract shapes come from a single participant. SLF: superior longitudinal fasciculus, UF: uncinate fasciculus.
